# *Solanum lycopersicum CLASS-II KNOX* genes regulate fruit anatomy via gibberellin-dependent and independent pathways

**DOI:** 10.1101/2022.07.06.498955

**Authors:** Amit Shtern, Alexandra Keren-Keiserman, Jean-Philippe Mauxion, Chihiro Furumizu, John Paul Alvarez, Ziva Amsellem, Naama Gil, Etel Motenko, Sharon Alkalai-Tuvia, Elazar Fallik, Nathalie Gonzalez, Alexander Goldshmidt

## Abstract

The pericarp is the predominant tissue determining the structural characteristics of most fruits. However, the molecular and genetic mechanisms controlling pericarp development remain only partially understood. Previous studies have identified that CLASS-II KNOX genes regulate fruit size, shape, and maturation in *Arabidopsis thaliana* and *Solanum lycopersicum*. Here we characterized the roles of the *Solanum lycopersicum* CLASS-II KNOX (TKN-II) genes in pericarp development via a detailed histological, anatomical, and karyotype analysis of the *TKN-II* knockdown (*35S:amiR-TKN-II*) fruits. We identify that *35S:amiR-TKN-II* pericarps contain more cells around their equatorial perimeter and fewer cell layers than the control. In addition, the cell sizes but not the ploidy levels of these pericarps were dramatically reduced.

Further, we demonstrate that fruit shape and pericarp layer number phenotypes of the *35S:amiR-TKN-II* fruits can be overridden by the *procera* mutant, known to induce a constitutive response to the plant hormone gibberellin. However, neither the *procera* mutation nor exogenous gibberellin application can fully rescue the reduced pericarp width and cell size phenotype of *35S:amiR-TKN-II* pericarps. Our findings establish that TKN-II genes regulate tomato fruit anatomy, acting via gibberellin to control fruit shape but utilizing a gibberellin-independent pathway to control the size of pericarp cells.

**Highlight:** Tomato *CLASS-II KNOX* genes regulate fruit and pericarp anatomy via GA-dependent and independent pathways.

## Introduction

Flowering plants (Angiosperm) dominate present-day terrestrial habitats and are the primary animal and human food source. The key distinctive feature of Angiosperm plants is the production of fruits, which facilitate seed protection and dispersal and display a vast diversity of shapes, sizes, and types (Clausing et al., 2000; Knapp, 2002; Pabon-Mora and Litt, 2011; Wang et al., 2015; Łangowski et al., 2016). In the majority of the Angiosperm species, the fruit anatomy is determined through the development of pericarp tissues derived from the ovary wall via differential cell divisions and cell expansion associated with endoreduplication (Cheniclet et al., 2005; Chevalier et al., 2011; Chevalier et al., 2014; Renaudin et al., 2017; van der Knaapa and Østergaard 2018; Ripoll et al., 2019; Dong and Østergaard 2019; Mauxion et al., 2021). The orientations and rates of cell divisions of the ovary/ pericarp cells were shown to control fruit shape (Wu et al., 2018; Lazzaro et al., 2018; van der Knaapa and Østergaard, 2018), while fruit weight and size correlate with the pericarp width determined by the number of cell layers and cell sizes (Cheniclet et al., 2005; Ripoll et al., 2019).

The size of the pericarp cells is tightly correlated with their ploidy levels (Cheniclet et al., 2005; Chevalier et al., 2011; Chevalier et al., 2014; Azzi et al., 2015; Renaudin et al., 2017; Ripoll et al., 2019; Musseau et al., 2020). However, it was demonstrated that the suppression of endoreduplication reduces cell ploidy but not always the pericarp cell size (Nafati et al., 2011), suggesting that endoreduplication alone does not determine cell size but instead acts as a limiting factor for cell expansion (Nafati et al., 2011; Chevalier et al., 2014).

Over the last three decades, several molecular factors that control pericarp cell division, endoreduplication, and cell expansion have been identified (Chevalier et al., 2014; Monforte et al., 2014; Azzi et al., 2015; van der Knaapa and Østergaard 2018; Dong and Østergaard 2019; Fenn and Giovannoni 2021; Mauxion et al., 2021). Different studies indicate that the plant hormones gibberellin (GA), auxin, and cytokinin (CK) are involved in the regulation of pericarp cell size and endoreduplication levels (Vivian-Smith and Koltunow, 1999; Marti et al., 2007; Su et al., 2014; Matsuo et al., 2012; Mauxion et al., 2021). However, the molecular networks through which the developmental and hormonal signals are integrated to control pericarp cell expansion remain poorly understood (Mauxion et al., 2021).

Previous studies in *Arabidopsis thaliana* and *Solanum lycopersicum* have identified CLASS-II KNOX genes controlling fruit size, shape, and maturation (Furumizu et al., 2015; Keren-Keiserman et al., 2022). In the current study, we used the previously described *35S:amiR-TKN-II* line shown to drive a ubiquitous knockdown of the SlCLASS-II KNOX (TKN-II) genes and phenocopy *slknatII3 slknatII7/+* mutants (Keren-Keiserman et al., 2022) to characterize the roles of TKN-II genes in the development of the *Solanum lycopersicum* (tomato) pericarps. We show that TKN-II genes regulate multiple aspects of pericarp development, including the orientation of cell divisions and the extent of cell expansion. We also show that the decreased number of pericarp cell layers and the altered fruit shape phenotypes of *35S:amiR-TKN-II* fruits can be overridden by the *procera* mutant, which is known to induce a constitutive response to the plant hormone gibberellin (GA) (Marti et al., 2007; Jasinski et al., 2008; Livne et al., 2015). However, neither the *procera* mutation nor ectopic GA application can fully complement the pericarp cell size phenotypes of the *TKN-II* knockdown fruits. These findings suggest that TKN-II genes utilize GA-dependent and independent pathways to regulate fruit and pericarp anatomy. This work and our recent report (Keren-Keiserman et al., 2022) advocate that TKN-II genes control multiple fruit traits by utilizing diverse molecular and hormonal networks at different stages of fruit development.

## Material and Methods

### Plant material

A commercial tomato M82 cultivar was used as a control. The transgenic *35S:amiR-TKN-II* line was previously described by Keren-Keiserman et al. (202_2_). The *procera*-loss-of-function allele was initially described by Jasinski et al. (2008). The allele’s introgression F4BC6 into the M82 background used in this study was described by Livne et al. (2015). The *35S:PROΔ17* plants were described by Nir et al. (2017). The double *35S:amiR-TKN-II pro* line was developed by crossing *pro* pollen on *35S:amiR-TKN-II* ovules, then applying *pro* pollen on the *35S:amiR-TKN-II /+ pro/+* F1 plants and screening for *35S:amiR-TKN-II /+ pro* (homozygous) plants in F1BC2 progeny. These plants were selfed (using available pollen) to recover about 30-50 seeds per plant. The F2BC2 plants were selfed again, then homozygous *35S:amiR-TKN-II pro* plants were identified based on the F3BC2 progeny segregation analysis.

### Plant Growth conditions

Plants were grown at ARO Volcani temperature-controlled greenhouse during the winter (September-April) and spring (March-June) seasons using the standard agronomic practice, under the average daily temperatures of 25-30^0^C and average night temperatures of 18-20^0^C. All described plants and phenotypes were grown, observed, and measured at least once during each season.

### Gibberellin treatment

ProGibb (GA3)(Valent BioScience Libertyville, Illinois, US) and 0.1% (by volume) of Triton™ X-100 (T8787, Sigma-Aldrich, St. Louis, Missouri, US) were used to prepare the hormonal solutions. The intact fruits were submerged in the solution for 30-40 seconds during each treatment. The treatment was initiated at 7 DAP fruits and applied once a week until the fruits reached the 35 DAP stage. At the same time, the control fruits were submerged in water with 0.1% Triton™ X100. The 40 DAP fruits were collected for measurements and imaging.

### Tissue preparation for histology, imaging, and image analysis

Reisin embedded histological ovary sections were prepared according to the protocol described by Izhaki et al. (2018). Ovary and pericarp tissues used for high-resolution fluorescent imaging were fixed overnight in 2.5% Paraformaldehyde solution, mounted in the Agarose LM (Thermo Fisher scientific H26417, Waltham, Massachusetts, US) frozen, sectioned using Leica SM2010R Sliding Microtome with a dry ice tray, and stained with Fluorescent Brightener 28 (F3543 Sigma-Aldrich, St. Louis, Missouri, US) and Propidium Iodide (P4864 Sigma-Aldrich, St. Louis, Missouri, US) or DAPI (D9542, Sigma-Aldrich, St. Louis, Missouri, US) according to the protocol described by Goldshmidt et al. (2008). The image acquisition was made using a Leica SP8 laser scanning microscope (Leica, Wetzlar, Germany), equipped with a solid-state laser with 405 and 552 nm light, PL APO 10x/0.4 (W.D.) 2.2mm, PL APO 20x/0.75 (W.D.) 0.62m or PL APO 40x/1.1 water (W.D.) 0.65mm objectives, and Leica Application Suite X software (LASX). Fluorescent brightener-28 and PI/RFP emission signals were detected with PMT and HyD (hybrid) detectors in ranges of 415–490 and 560–660 nm, respectively. Following the imaging, all confocal images were procced with Imaris x 64 8.1.2 (Bitplane A.G, Belfast, UK) to obtain 3D reconstructions and 2D image frames. Fresh ovaries, immature fruits, and pericarp tissues used for cell layer count, ovary/fruit perimeter, visual cell size assessment, and width measurements were cut using the razor blade, dipped into 0.5% Toluidine Blue O (TBO) (T3260, Sigma-Aldrich, St. Louis, Missouri, US) water solution, washed twice in water, and immediately imaged using the Nicon SZM25 stereoscope; SHR Plan Apo 1X WD60 objective. The fully grown and mature fruits were cut and imaged using a Canon Power Shot SX520 HS camera. Fiji –Image J 64 and CellSeT (Pound et al., 2012) software were used to take the measurements following the imaging and image processing.

### Transcription levels assessment by the quantitative PCR test

All plant tissues used to assess the transcription levels were collected into liquid nitrogen 3-4 hours after dawn. The RNA was extracted using the Gene All, “RiboEx – total RNA isolation” kit (#301-001); the” DNA-free DNase Treatment & Removal” kit (AM1906 Invitrogen, ThermoFisher Scientific, Waltham, Massachusetts, US) was used to remove any DNA contaminations. Subsequently, one μg of RNA was used to prepare cDNA with the “High capacity RNA–to-cDNA” kit (#4387406 Applied biosystems-ThermoFisher Scientific, Waltham, Massachusetts, US). The qPCR analysis of the total cDNA was performed using the “Fast SYBR Green Master Mix “(# 4385612 Applied biosystems-ThermoFisher Scientific, Waltham, Massachusetts, US), gene-specific primers, and SAND primers (Expósito-Rodríguez et al., 2008) as a control (Suppl. **Tabel 1**), the Applied biosystems-ThermoFisher Scientific, Waltham, Massachusetts, US -Step One Plus Real-time PCR system and software. The Relative expression (RQ) values were calculated as described (Pfaffl, 2004).

### Calculations and statistical analysis

All plants were randomized in the greenhouses. Only the plants growing in the same season were compared to the controls growing under the same conditions. Estimations of the epidermal/E1 cell numbers across fruit /ovary equatorial perimeters were performed by dividing fruit/ovary perimeters by the average length of the epidermal/E1 cells. The means differences were compared by the Student’s –t-test followed by the Benjamini–Hochberg procedure to control for the False Discovery Rates (Benjamini and Hochberg 1995) using the R-language, “tydiverse,” and “rstatix” packages.

### Ploidy level analysis

Frozen fruit pericarps were chopped in 200 μl of chilled CyStain UV Precise P Nuclei Extraction Buffer (Sysmex, Kobe, Japan) using a razor blade. The suspension was filtered through a 50-μm nylon filter, and 800 μl of chilled CyStain UV Precise P Staining Buffer (Sysmex, Kobe, Japan) was added to the isolated nuclei. The 2,000–10,000 nuclei DNA content was measured using a CyFlow Space flow cytometer (Sysmex, Kobe, Japan) and analyzed with FloMax software (Sysmex, Kobe, Japan).

## Results

### SlCLASS-II KNOX genes regulate pericarp anatomy

Recent examination of the TKN-II genes roles in tomato fruit development has shown that the *tknII3, tknII5*, and *tknII7* single mutants significantly reduce fruit size and shape. These phenotypes become even more dramatic in the *tknII3 tknII7/+* and *35S:amiR-TKN-II* fruits. The visual assessment of *tknII3 tknII7/+* and *35S:amiR-TKN-II* fruits suggests that their reduced size is associated with the visible reduction of their pericarps thickness (Keren-Keiserman et al., 2022) (Fig.1A). In the current study, we used the *35S:amiR-TKN-II* line shown to significantly down-regulate transcription of all TKN-II clade members and phenocopy *tknII3 tknII7/+* line (Keren-Keiserman et al., 2022) to characterize the roles of TKN-II genes in the regulation of pericarp development and fruit anatomy. To decipher at what stage of fruit development TKN-II activity is responsible for the phenotypic differences observed between the mature M82 and *TKN-II knockdown* pericarps (Fig.1A), we compared pericarp development of M82 and *35S:amiR-TKN-II* fruits from the anthesis stage (Fig.1B) to fully grown fruits at 40 Days After Pollination (DAP) stage (Fig.1A). First, we used stereoscope and confocal images of the ovary wall and pericarp sections (Fig.1A-B) to count the number of abaxial-adaxial cell layers (Fig. 1C), to measure the equatorial ovary/pericarp perimeters and average lateral length of the top exocarp layer (E1) cells (Renaudin et al., 2017), and to estimate the number of E1 cells around the equatorial perimeter (Fig. S1A-F;1D).

**Figure 1:**
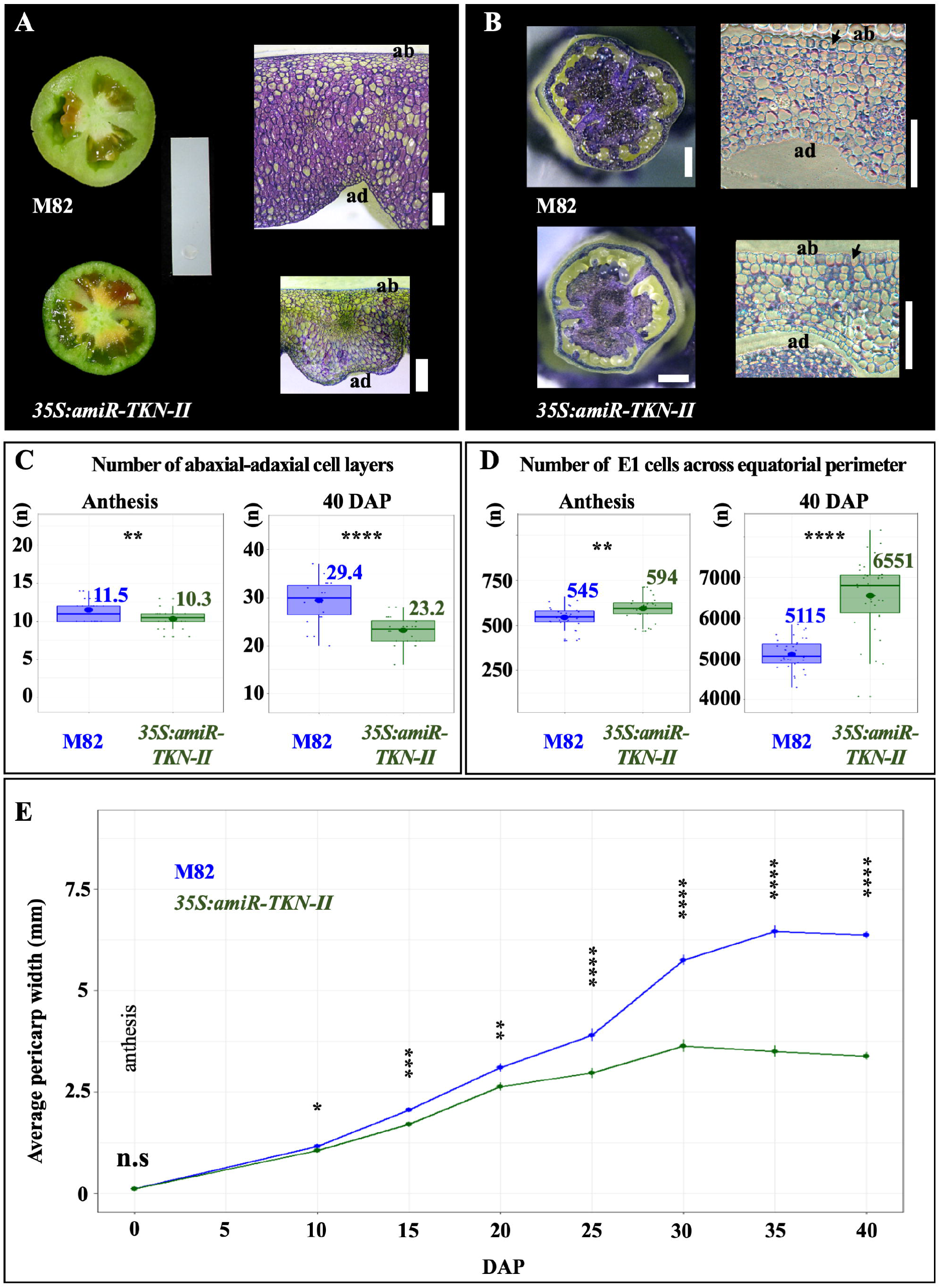
Knockdown of the *SlCLASS-II KNOX* genes alters pericarp anatomy. **A:** M82 and *35S:amiR-TKN-II* fresh-cut equatorial sections of 40 Days After Pollination (DAP) fruits (left column; size bar is 6cm) were used to assess pericarp thickness, measure the equatorial pericarp perimeter, and assess the number of E1-layer cells across the fruit equator. Stereoscope images of the Toluidine blue (TB) stained fresh-cut pericarps from similar 40 DAP fruits (right column; size bars are 1mm) were used to count the number of the pericarp abaxial-adaxial cell layers and measure pericarp thickness. **B:** M82 and *35S:amiR-TKN-II* fresh (left column; size bars are 1mm) and resin embedded (right column; size bars are 100μm) equatorial sections of anthesis ovaries stained with TB were used to count the abaxial-adaxial cell layers, measure ovary perimeter, and ovary wall width, and estimate the numbers of E1-layer cells across ovary wall equator. **C:** Box plots displaying the distributions and average numbers (large dot and number next to box plot) of the abaxial-adaxial cell layers in M82 (blue) and *35S:amiR-TKN-II* (dark green) anthesis ovaries (left panel), and 40 DAP pericarps (right panel). Small dots represent individual measurements. **D:** Box plots displaying the estimated distributions and average numbers (large dot and number next to box plot) of the E1 cells across the equatorial perimeters of the M82 (blue) and *35S:amiR-TKN-II* (dark green) anthesis ovaries (left panel), and 40 DAP pericarps (right panel). Small dots represent individual measurements. **E:** Line plots representing the M82 (blue) and *35S:amiR-TKN-II* (dark-green) average (round-dots) equatorial pericarp expansion during anthesis (0 DAP) to 40 DAP stages. The error bars represent standard errors. In **C-E**: Student’s-t-test H_0_ probabilities are presented as (**n.s**) - p.val > 0.05; (**\***)-p.val≤ 0.05; (**\*\***) - p.val≤ 0.01; (**\*\*\***) - p.val≤ 0.001; (****) - p.val≤ 0.0001. In **C-D**: the number of sampled fruits and ovaries for each stage and genotype is 30. In **E**, the number of sampled fruits/ovaries for each stage and genotype is ≥25.

We found that *35S:amiR-TKN-II* anthesis-stage ovary wall and fully grown 40 DAP pericarps had fewer abaxial-adaxial cell layers than the control (Fig. 1C). At anthesis, the average lateral length of the transgenic E1 ovary wall cells along the equatorial fruit perimeter was similar to that of M82 cells from the E1 layer (Fig.S1C). However, the equatorial perimeter of the *35S:amiR-TKN-II* ovaries was significantly greater than the control, presumably due to a greater number of E1 cells in *35S:amiR-TKN-II* lines (Fig.S1A) (Fig. 1D-left panel).

The equatorial perimeters of the fully-grown pericarps and the average lateral length of the E1 cells around them were significantly reduced in *35S:amiR-TKN-II* fruits compared to M82 fruits (Fig.S1D-F). The number of E1 cells around the 40 DAP fruit equatorial perimeter was estimated to be about 1400 more in the *35S:amiR-TKN-II* pericarps than in M82 pericarps (Fig. 1D-right panel). These observations suggest that the knockdown of the TKN-II genes triggered a reduction in the number of pericarp cell layers but led to an increased number of E1 cells along the equatorial fruit perimeter.

At the anthesis-stage, *35S:amiR-TKN-II* ovary wall had about one abaxial-adaxial cell layer less than the M82 ovary wall (Fig.1C-left panel), which did not lead to significant differences in the ovary wall width at this stage (Fig. 1E). At 10 DAP, *35S:amiR-TKN-II* pericarps had, on average, six-cell layers less than the M82 control (Fig.S2), resulting in a small but significant reduction in the average pericarp width (Fig. 1E). Following the 10 DAP stage, no additional pericarp layers were formed in both lines (Fig.S2); however, the pericarp width differences between the genotypes increased, leading to significantly thinner pericarps in fully grown *35S:amiR-TKN-II* fruits (Fig.1A;1D). These observations imply that downregulation of TKN-II transcription impairs the expansion of the pericarp cells.

### *SlCLASS-II KNOX* activity regulates the size of tomato pericarp cells

To further assess the effect of TKN-II knockdown on the pericarp cell expansion, we used high-resolution confocal images of 10, 20, and 40 DAP pericarp sections (Fig. S3) and Cell SeT software (Pound et al., 2012) to survey the distributions of the cell area sizes in the pericarps of M82 and *35S:amiR-TKN-II* fruits. At 10 DAP, pericarp tissues of both genotypes had equally-sized cells (Fig. 2A-top panel). This observation suggests that, at this stage, the reduction in the total pericarp width observed in *35S:amiR-TKN-II* fruits (Fig. 1E) was associated with decreased pericarp cell layer numbers (Fig. S2). At 20 DAP, a lower proportion of the 0.01-0.05 mm2 (large) cells was detected in *35S:amiR-TKN-II* pericarps (Fig. 2A-middle panel). Finally, the fully grown (40 DAP) pericarps of the *35S:amiR-TKN-II* fruits were found to have significantly larger proportions of small cells (≤0.01mm^2^) and a significantly smaller proportion of large (0.01-0.05mm^2^) and giant (0.05-0.1mm^2^) cells than the controls (Fig. 2A - bottom panel). In the pericarps of both genotypes, a similar low percent of the extra-giant cells (≥ 0.1mm^2^) was found, and cells larger than 0.13mm^2^ were detected only in the M82 pericarps (Fig. 2A-bottom panel).

**Figure 2.**
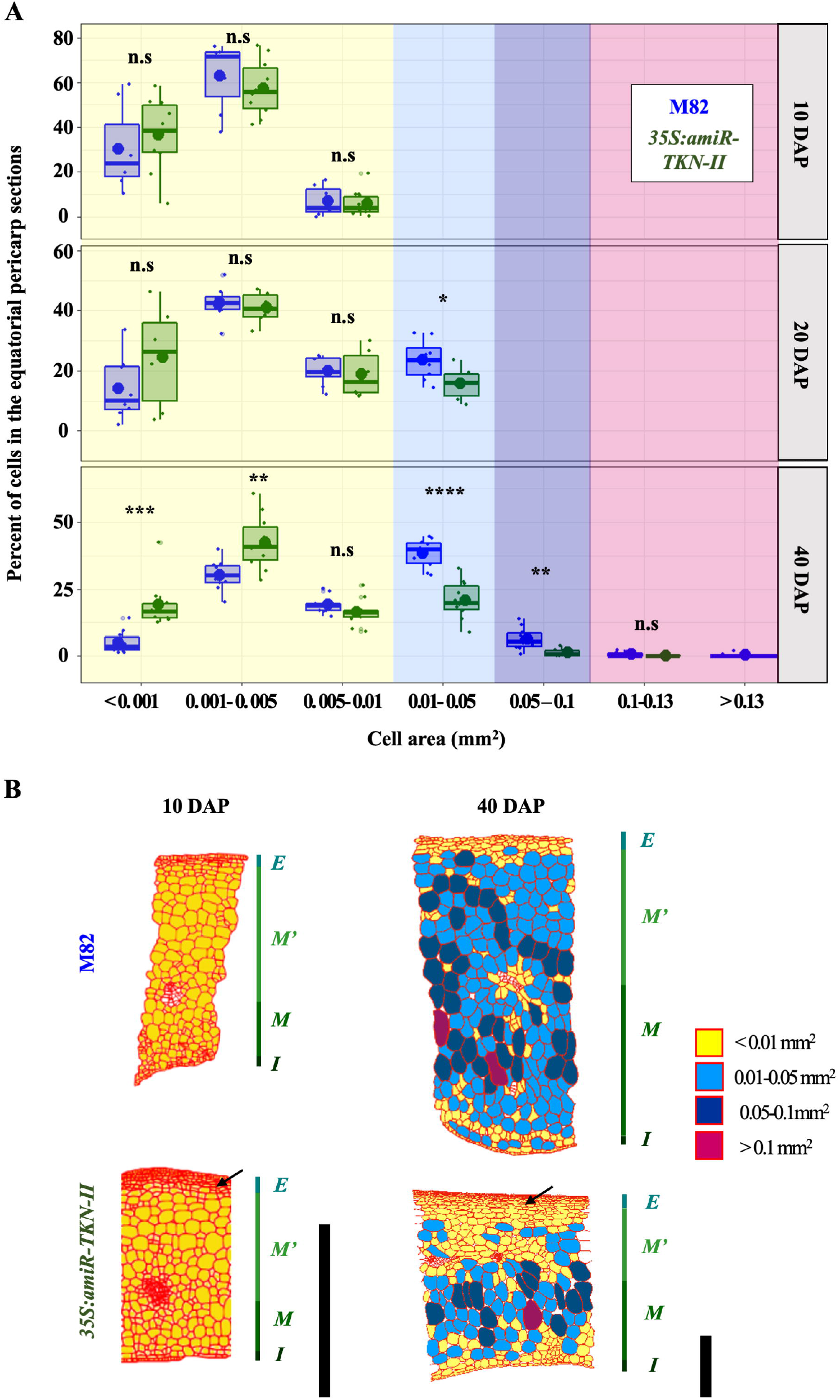
Knockdown of the *SlCLASS-II KNOX* genes reduces the proportion of large and giant cells and increases the proportion of small cells within the pericarp. **A:** Box-plots displaying the distribution and average percent (large dots) of different size range cells in the pericarps of M82 (blue) and *35S:amiR-TKN-II* (dark-green) at 10 DAP (top panel), 20 DAP (middle panel), and 40 DAP (bottom panel). **Y-scale** indicates the percentage of cells in the pericarp sections, **X-scale** indicates cell area size ranges in mm^2^. Background colors of each panel indicate the cell area ranges presented in **B**. Small dots represent individual measurements. Student’s-t-test H_0_ probabilities are presented as (**n.s**) - p.val > 0.05; (*)- p.val≤ 0.05; (**) - p.val≤ 0.01; (***) - p.val≤ 0.001; (****) - p.val≤ 0.0001. The number of sampled pericarps for each stage and genotype ≥ 7. **B:** Pericarp cartograms generated by CellSeT software (Paound et al., 2012), representing the distributions of cell size-types at the 10 and 40 DAP- M82 (top) and *35S:amiR-TKN-II* (bottom) equatorial pericarp sections. Cell colors indicate the range of the cell area size: **yellow** – cells smaller than 0.01mm^2^ (small cells); **light blue**- cells larger than 0.01mm^2^ but smaller than 0.05 mm^2^ (large cells); **dark blue-** cells larger than 0.05mm^2^ but smaller than 0.1 mm^2^ (giant cells); **magenta**- cells larger than 0.1mm^2^ (extra giant cells). The color scale next to each cartogram and letters mark the pericarp layers and tissue types according to (Renaudin et al., 2017): **marine-green** (E) – exocarp; **forest-green** (M’) –upper mesocarp (above the vein); **dark-green** (M)-low mesocarp (below the vein). **Deep-dark green** (I) – endocarp. The size bars are 1mm.

Following the quantitative analysis of the cell size-types distribution, we used Cell SeT (Pound et al., 2012) generated cartograms (Fig. 2B) of high-resolution confocal images (Fig. S3A-C) to assess the spatial distributions of cell sizes in the pericarp layers of the two genotypes. The pericarp layers were defined and named according to previously published nomenclature (Renaudin et al., 2017). At 10 DAP, a larger amount of the longitudinally oriented cell layers were detected in the *35S:amiR-TKN-II* exocarp (E) – upper mesocarp (M’) boundary region compared to M82 pericarps of the same age (Fig. 2B-left panel marked with an arrow; Fig.S3A marked with an arrow in the right panel and shown in higher magnification in the inset image at the bottom). A similar trend was observed at the fully grown (40 DAP) pericarps of these genotypes (Fig. 2B-right panel marked with an arrow; Fig. S1E; S3C- right panel marked with an arrow). The majority of the M82 upper mesocarp (M’) cells belonged to the 0.01-0.05mm2 (large cell) size group, whereas most of the *35S:amiR-TKN-II* M’ cells were smaller than 0.01mm^2^ (Fig.2B-right panel marked with arrow). The low-mesocarp layers (M) of both genotypes had large and giant cells; however, the transgenic M regions had a greater proportion of the smaller-sized cells than the control (Fig. 2B-right panel).

Together, these observations demonstrate that TKN-II genes regulate the size of tomato pericarp cells.

### *SlCLASS-II KNOX* genes impact the dynamics of pericarp endo-reduplication

Multiple studies of plant tissues demonstrate that cell size is tightly and positively correlated to ploidy level (Chevalier et al., 2011; Azzi et al., 2015). Upon establishing that TKN-II genes control the pericarp cell size (Fig. 2A-B), we compared ploidy levels of *35S:amiR-TKN-II* and M82 developing pericarps. As a control, we assessed the ploidy levels of mature leaves, which displayed similar ploidy distributions in both genotypes, with the majority of the cells being at the 2C level (Fig. 3 -top panel). Similar to the mature leaves, no differences in the cell-ploidy distributions between the genotypes were found in the 10 DAP pericarp tissues. At the 10 DAP stage, 2C cells represented 25% of the total cell number. The pericarp cell ploidy increased with fruit development in both genotypes, and no 2C cells could be detected after the 10 DAP stage. At the 20 DAP stage, a lower proportion of the 32C cells was detected in *35S:amiR-TKN-II* pericarps. Subsequently, at the 25 DAP stage, a higher proportion of the 4C cells and lower proportions of the 16C and 32C cells were detected in the *35S:amiR-TKN-II* pericarps. A similar trend was observed at the 30 DAP stage, where lower proportions of the 16C, 32C, and 64C cells were found in the pericarps of the *35S:amiR-TKN-II* fruits (Fig. 3). Surprisingly in fully grown 40 DAP pericarps, the ploidy distribution differences between the two genotypes appeared less dramatic than their cell size differences. While at the 40 DAP *35S:amiR-TKN-II* pericarps, a significantly higher proportion of small cells and lower proportions of the large and the giant cells were detected than in 40 DAP M82 pericarps (Fig. 2A-bottom panel), the comparison of the ploidy distributions of similar tissues identified only a higher number of the 8C and a lower number of 256C cells in the transgenic pericarps (Fig. 3 - 40DAP panel). These results suggest that reduced TKN-II transcript levels cause a delay in the progression of endoreduplication during fruit development that is partially compensated at the end of fruit growth.

**Figure 3:**
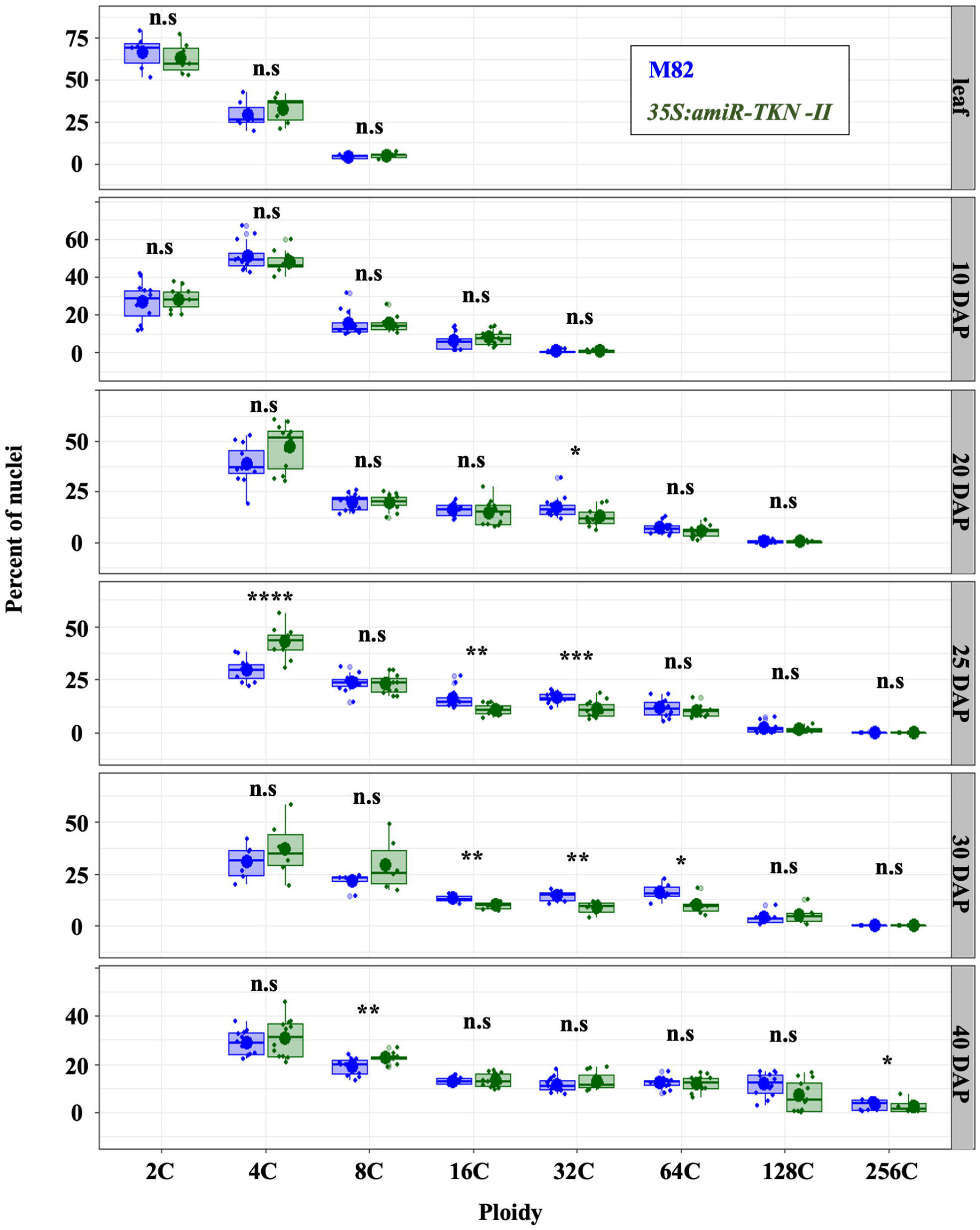
Downregulation of the *TKN-II* transcription reduces the proportions of high ploidy nuclei in 20,25,30 and 40 DAP pericarps. **A:** Box-plots dispalying the distribution and average percent (large dots) of different ploity nuclei at M82 (blue) and *35S:amiR-TKN-II* (dark green) fifth leaf, 10, 20, 25, 30 and 40 DAP equatorial pericarp tissues (horizontal panels). **X-scale** indicates nuclei ploidy levels; **Y-scale** percent of the nuclei in the sampled tissue. Small dots represent individual measurements. Student’s-t-test H_0_ probabilities are presented as (**n.s. – not significant**) (p-value)> 0.05; (**\***);(p-value) ≤ 0.05; (**\*\***) (p-value) ≤ 0.01; (**\*\*\***) (p-value) ≤ 0.001; (**\*\*\*\***) (p-value) ≤ 0.0001.The number of sampled fruits at each stage for each genotype is ≥ 5. The number of sampled leaves = 6.

### The *procera* mutant compensates for *35S:amiR-TKN-II* leaf and fruit shape phenotypes

The analysis of the developing M82 and *35S:amiR-TKN-II* pericarps revealed that reducing the *TKN-II* transcription levels inhibits the pericarp expansion after the 10 DAP stage (Fig. **1E**; **2A-B**). Previous research has established that the plant hormone gibberellin (GA) promotes the expansion of the pericarp cells (Bonger-Kibler and Bangerth, 1982; Marti et al., 2007; Shinozaki et al., 2018). Moreover, it was demonstrated that the GA activity mediates leaf phenotypes induced by the ectopic expression of the *SlCLASS-I KNOX* genes [antagonists of *SlCLASS-II KNOX* genes (Furumizu et al., 2015)] (Jasinski et al., 2008). Based on these findings, we wished to test whether the inhibition of pericarp expansion in *35S:amiR-TKN-II* fruits is associated with GA.

First, we determined the transcription levels of all pericarp expressed GA metabolic (GA20OX) and catabolic (GA2OX) (Hedden 2020) genes, as well as the expression of the GA response regulatory gene PROCERA (PRO) (Marti et al., 2007: Jasinski et al., 2008; Livne et al., 2015) (Fig. S4A) in the developing M82 and *35S:amiR-TKN-II* pericarps.

The only gene in which we found a significant change in expression level was GA catabolic gene *SlGA2OX2*. The levels of the *SlGA2OX2* were significantly upregulated in 25-40 DAP pericarps of the *35S:amiR-TKN-II* fruits (Fig. S4B), coincident with the significant differences in pericarp development (Figure 1A-E; Fig. S4B-middle panel).

Based on these results, we hypothesized that *35S:amiR-TKN-II* fruit phenotypes are a consequence of reduced GA activity and could be restored by constitutive activation of the GA response.

Previous studies have established that *Solanum lycopersicum procera* (*pro*) mutation induces a constitutive response to GA across all plant tissues (Marti et al., 2007; Jasinski et al., 2008; Livne et al., 2015). We introgressed the *35S:amiR-TKN-II* transgene into the *pro* mutant background to examine whether it can rescue *TKN-II* knockdown phenotypes. The *35S:amiR-TKN-II* leaves are highly complex and dark-green colored (Keren-Keiserman et al., 2022; Fig. S5A), whereas the *pro* mutant leaves, grown under the same conditions, are light green and have less complex shapes than the M82 control (Jasinski et al., 2008; Livne et al., 2015; Fig. S5A). We detected that the leaves of the *35S:amiR-TKN-II pro* plants had shape patterns similar to *pro* mutant leaves. However, the color of these leaves remained dark-green as in *35S:amiR-TKN-II* leaves (Fig.S5A).

Next, we measured the ovary shape indexes [ratio between the ovary length and maximal ovary width (Brewer et al., 2006)] at the anthesis and the 40 DAP stages of M82, *pro, 35S:amiR-TKN-CL-II*, and *35S:amiR-TKN-II pro* lines. Analysis of shape index in anthesis stage detected significant differences between the M82, *35S:amiR-TKN-II*, and *pro* genotypes M82 ovaries had a shape index of 1.06 corresponding to a slightly elongated ellipsoid shape, *35S:amiR-TKN-II* ovaries had an average shape index of 0.96 corresponding to an oblate ellipsoid shape, and *pro* ovaries had an average shape index of 1.14 corresponding to a prolate ellipsoid shape. The average shape index of the *35S:amiR-TKN-II pro* ovaries was similar to the shape index of the *pro* mutant ovaries (Fig. 4A-B). The shape indexes of the mature fruits from the four genotypes were similar to those measured at the anthesis stage ovaries (Fig. 4C). The mature fruit shape differences between the M82, *35S:amiR-TKN-CL-II*, and *pro* lines could be visually detected, whereas the shapes of *35S:amiR-TKN-II pro* and *pro* fruits were similar (Fig. 4D). Interestingly, the mature fruits of the *35S:PROΔ17* line, shown to repress the GA response (Nir et al., 2017), develop oblate-shape fruits similar to the *35S:amiR-TKN-II* fruits (Fig. S5B). These findings suggest that constitutive GA response can override *35S:amiR-TKN-II* leaf and ovary/fruit shape phenotypes established at the anthesis stage (Fig. 4A-B).

**Figure 4:**
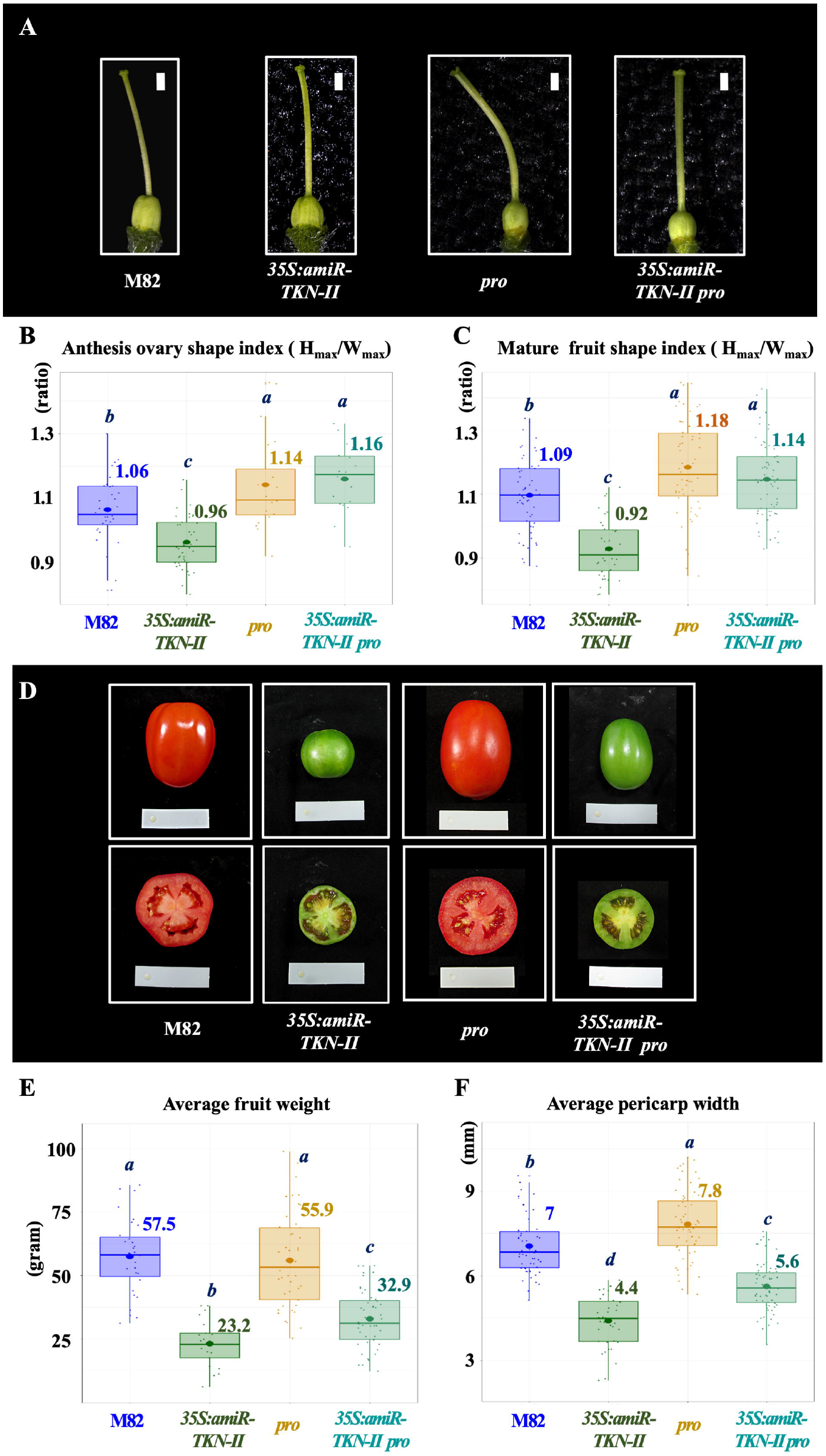
*procera (pro)* loss-of-function mutation has an epistatic effect on *35S:amiR-TKN-II* fruit shape and partially complements its pericarp width. **A:** Stereoscope images of the dissected anthesis stage ovaries from M82, *35S:amiR-TKN-II, pro*, and *35S:amiR-TKN-II pro* lines (shown in individual panels) were used to computationally measure ovary shape indexes [the ratio of ovary length to ovary width (Brewer et al., 2006) (**B**)] of the respective genotypes. The size bar of each panel = 1mm. **B:** Box plots displaying the distributions and averages (large dots and numbers next to each box) of the M82, *35S:amiR-TKN-II, pro*, and *35S:amiR-TKN-II pro* ovary shape indexes. **C:** Box plots displaying the distributions and averages (large dots and numbers next to each box) of the M82, *35S:amiR-TKN-II, pro*, and *35S:amiR-TKN-II pro* mature fruit (**D**) shape indexes [measured as the ratio of fruit length to its maximum width (Brewer et al., 2006)]. **D:** Whole fruit (top row panels) and equatorial sections (bottom row panels) of the 50 DAP M82, *35S:amiR-TKN-II, pro*, and *35S:amiR-TKN-II pro* fruits. In each panel the size bar = 6cm. **E:** Box plots displaying the distributions and the average weight (large dots and numbers next to each box) of the 40 DAP M82, *35S:amiR-TKN-II, pro*, and *35S:amiR-TKN-II pro* fruits. **F:** Box plots displaying the distributions and the average pericarp width (large dots and numbers next to each box) of the mature M82, *35S:amiR-TKN-II, pro*, and *35S:amiR-TKN-II pro* fruits. In **B-C; E-F:** Box and dot colors represent values of the following genotypes: M82-blue, *35S:amiR-TKN-II –*dark green, *pro-* marigold, *35S:amiR-TKN-II pro*-marine-green. Small dots represent individual measurements. Different letters indicate Student’s-t-test (adjusted using Benjamini–Hochberg method) H_0_ probability values ≤ 0.05, the similar letters indicate probability values > 0.05. In **B**, the number of measured ovaries for each genotype is ≥ 18; In **C** and **E**, the number of measured fruits for each genotype is ≥ 39; In **F**, the number of fruits measured for each genotype is ≥ 21.

### The *procera* mutant does not fully complement pericarp width and cell-size phenotypes of *35S:amiR-TKN-II* fruits

The *pro* mutants were shown to increase pericarp width and cell size (Marti et al., 2007; Shinozaki et al., 2018), which are decreased in the *35S:amiR-TKN-II* fruits (Fig. 1A;1E; 2A-B; 4D). To test whether the constitutive GA response induced by the *pro* mutation can rescue *35S:amiR-TKN-II* pericarp phenotypes, we examined the M82, *35S:amiR-TKN-II, pro*, and *35S:amiR-TKN-II pro* mature fruits.

Previous reports have shown that *pro* mutants may develop facultative fruit parthenocarpy (Marti et al., 2007; Livne et al., 2015; Shinozaki et al., 2018). Parthenocarpy and insufficient fruit pollination were demonstrated to reduce fruit size and pericarp cell expansion (Marti et al., 2007; Shinozaki et al., 2018; Ripoll et al., 2019). In order to correctly assess the effects of the *pro* - *35S:amiR-TKN-II* genetic interaction on the fruit and pericarp development and to control for the timing of the fruit set, the homozygous M82, *pro, 35S:amiR-TKN -II*, and *35S:amiR-TKN-II pro* anthesis stage ovaries were hand-pollinated using the M82 pollen. At 40 DAP, the marked hand-pollinated fruits were collected for measurements. A subset of the fruits was imaged at 50 DAP when the M82 and *pro* fruits reached the Ripen Red (RR) stage, but *35S:amiR-TKN-II* and *35S:amiR-TKN-II pro* fruit pericarps remained green (Fig. 4D).

First, we compared the average 40 DAP fruit weight of the four genotypes. No significant weight differences between the M82 and *pro* fruits were identified. The *35S:amiR-TKN-II* fruits were significantly lighter and smaller than all the other genotypes, whereas *35S:amiR-TKN-II pro* fruits were heavier than the *35S:amiR-TKN-II* fruits but still significantly lighter and smaller than M82 and *pro* fruits (Fig.4E). Next, we measured M82, *pro, 35S:amiR-TKN-CL-II*, and *35S:amiR-TKN-II pro* pericarp width at 40 DAP. We found that the pericarps of the *pro* mutant were thicker than M82 pericarps, which were thicker than the pericarps of the *35S:amiR-TKN-II pro* fruits. The pericarps of the *35S:amiR-TKN-II* fruits were the thinnest among the four genotypes (Fig. 4D-bottom panels; 4F). These observations suggest that *pro* mutation was insufficient to fully complement the pericarp width and weight phenotypes of the *35S:amiR-TKN-II* fruits.

To characterize pericarp width differences between the four genotypes at higher resolution, we first monitored cell numbers in the equatorial ovary sections at the anthesis stage (Fig. S6A). This assay confirmed the reduction of one ovary wall abaxial-adaxial cell layer in *35S:amiR-TKN-II* ovaries (Fig. 1B-C; Fig. S6A-B). No significant differences in the abaxial–adaxial ovary wall layers were identified between the rest of the genotypes (Fig. S6A-B). However, when in comparison to M82 ovaries the *35S:amiR-TKN-II* ovaries showed a significant increase in the number of E1 cells along their equatorial perimeter (Fig. 1B; 1D; Fig. S6C), the *pro* and *35S:amiR-TKN-II pro* ovaries had significantly less E1 cells across their equatorial perimeter than M82 ovaries (Fig. S6C). Based on the visual assessment of ovary sections (Fig. S6A) and the computational measurements, no significant differences in the ovary wall E1 cell lateral length and size were detected between the genotypes at the anthesis stage (Fig. S6A; S6D).

Next, we analyzed the 40 DAP equatorial pericarp sections of the four genotypes (Fig 5A). The 40 DAP pericarp cell layer counts detected a significant decrease in the pericarp cell layers and an increase (in comparison to M82) of E1 cells along the *35S:amiR-TKN-II* equatorial fruit perimeter (Fig.1C-D-right panel; 5B-C). The cell counts of the 40 DAP pericarps did not detect significant differences in the abaxial-adaxial layer numbers between the M82 and *pro* mutant or between the M82 and *35S:amiR-TKN-II pro* double mutant. However, *35S:amiR-TKN-II pro* fruits appeared to have fewer pericarp abaxial-adaxial layers than *pro* fruits (Fig. 5B). The mature fruits of the four genotypes maintained similar differences in the numbers of E1 cells across their equatorial perimeters as in their anthesis stage ovaries (Fig. S6C; Fig. 5C).

**Figure 5:**
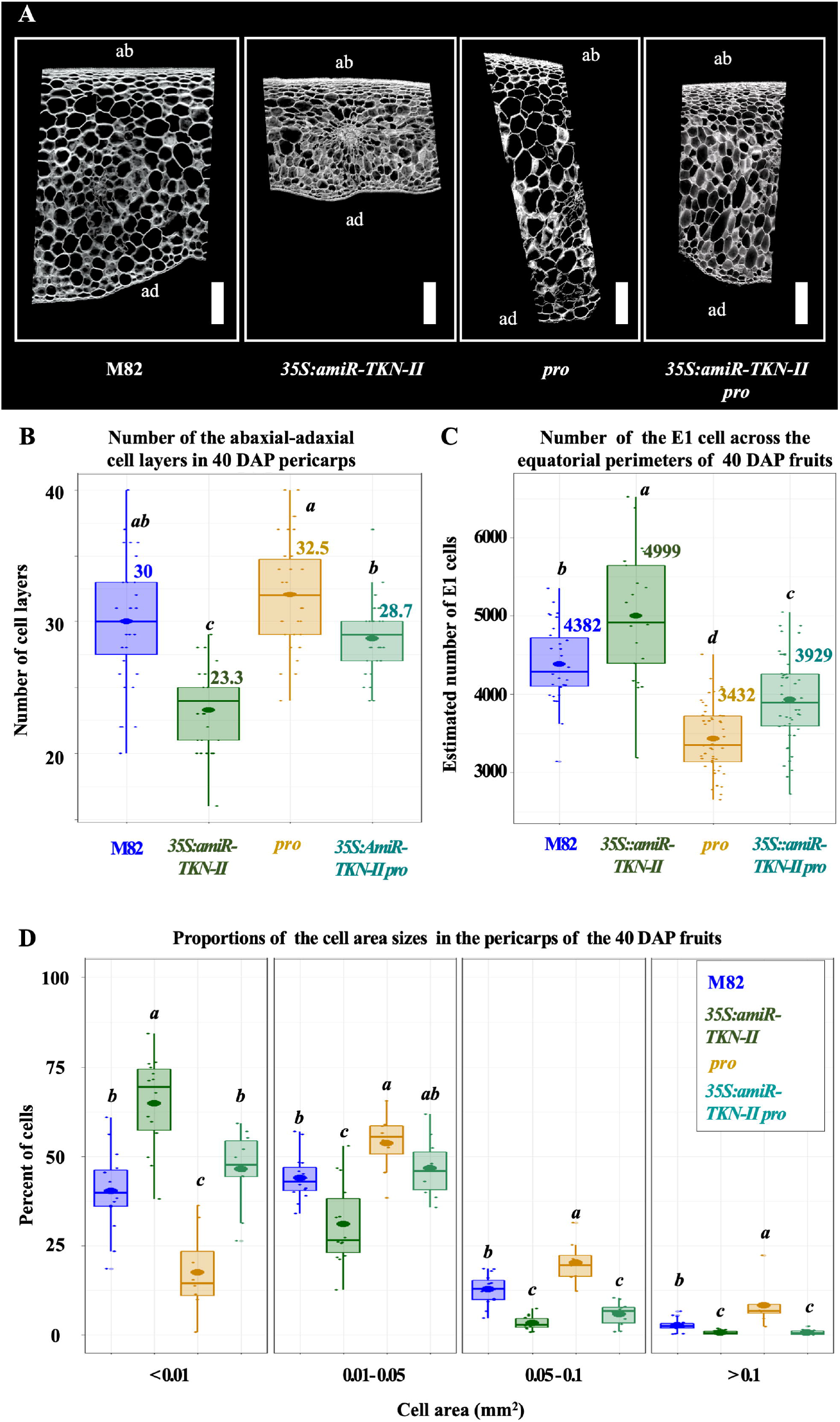
Histological analysis of the 40 DAP M82, *35S:amiR-TKN-CL-II, pro*, and *35S:amiR-TKN-II / pro* equatorial pericarp sections. (**A**) High-resolution Fluroscent Brightner-28 stained equatorial pericarp confocal images of the four genotypes presented in separate panels. The size bar of each panel = 1mm. The images were used to count the abaxial-adaxial cell layers [the abaxial pericarp side marked in as (**ab)**, the adaxial pericarp side marked in as an (**ad**)]. Distributions of the cell-layer numbers are presented in the box plot (**B**). Same images (**A**) were used to measure the cell area distributions within the pericarps (**D**). The number of the E1 cells (marked in **A** with arrows) across the equatorial fruit perimeter displayed at the box-plot chart (**C**) was estimated using the whole fruit equatorial sections. In **B-C**: Numbers next to each box-plot represent the mean cell-layer/ cell number of the respective genotype. In **B**, the number of sampled pericarp sections from different fruits is 30 per genotype; in **C**, the number of measured fruits is 30 per genotype **D:** Bar plots display the distributions of the cell area sizes measured by the CellSeT software (Paound et al., 2012) in the pericarps of M82 (blue) and *35S:amiR-TKN-II* (dark-green), *pro* (marigold), and *35S:amiR-TKN-II pro* (marine-green) at 40 DAP pericarps. **Y-scale** indicates the percent of the cells in the pericarp sections, **X-scale** indicates cell area size ranges in mm^2^. The number of the sampled pericarp sections is ≥ 10. In **B-D:** Small dots represent individual measurements. Different letters indicate Student’s-t-test H_0_ probability values (adjusted using Benjamini–Hochberg method) ≤ 0.05, the similar letters indicate probability values > 0.05. The means comparison was performed separately for each cell area group.

Next, we used the Cell SeT application (Pound et al., 2012) to assess the cell area distributions at the 40 DAP pericarps of the: M82, *35S:amiR-TKN-II, pro*, and *35S:amiR-TKN-II pro* fruits. We found that the fully grown *pro* pericarps had a smaller proportion of the small cells (<0.01mm^2^) and larger proportions of large (0.01-0.05mm^2^), giant (0.05-0.1mm^2^), and super-giant cells (>0.1mm^2^) than the rest of the genotypes. The comparable pericarps of *35S:amiR-TKN-II* fruits had a larger proportion of the small cells (<0.01mm^2^) and a smaller proportion of the large cells (0.01-0.05mm^2^) than the M82 pericarps. The proportion of small (<0.01mm^2^) cells and large (0.01-0.05mm^2^) cells in the *35S:amiR-TKN-II pro* pericarps were not significantly different from the control (M82). However, both the *35S:amiR-TKN-II* and *35S:amiR-TKN-II pro* pericarps had significantly smaller proportions of giant (0.05-0.1mm^2^) and super-giant (>0.1mm^2^) cells than M82 and *pro* pericarps of the same stage. (Fig. 5D).

The histological ovary/fruit analysis of the four genotypes suggests that *pro* mutation fails to fully complement the pericarp cell size phenotypes of the *TKN-II* knockdown fruits. To further examine the effect of *pro* mutation on the pericarp cells, we monitored 40 DAP pericarp ploidy levels of the four genotypes. The fully grown *pro* pericarps had a significantly lower percentage of the 4C and 8C nuclei and a significantly higher percent of the 128C and 256C nuclei than the other three genotypes. In addition, only in *pro* pericarp samples, 512C nuclei were detected (Fig. S7 marked with arrow). However, except for the higher percentage of 16C cells, the ploidy patterns of *35S:amiR-TKN-II pro* pericarps were similar to the ploidy patterns of the *35S:amiR-TKN-II* pericarps (Fig. S7).

Thus the pericarp ploidy analysis supports the rest of the observations (Fig. 4D; 4F; 5A; 5D; S7), showing that *35S:amiR-TKN-II* pericarp width and cell size cannot be fully complemented by the *procera* mutant.

### Fruit-specific GA_3_ application does not compensate for width differences between the M82 and *35S:amiR-TKN-II* pericarps

Previous reports indicate that phenotypic effects of *pro* mutation may depend on the relative strength of the specific *pro* allele (Shinozaki et al., 2018). To test whether the pericarp width of the *35S:amiR-TKN-II* fruits could be rescued by high concentrations of ectopically supplied GA, we applied four concentrations of 20, 40, 80, and 160 ppm of GA_3_ to the developing *35S:amiR-TKN-II* and M82 fruits.

The fruits of both genotypes responded to GA_3_ treatment by increasing the pericarp width and cell size (Fig. 6A-B; Fig. S8A). In both genotypes, the maximum response to hormone treatment was at 40 ppm, resulting in a significant increase of pericarp width by 2mm and plateauing at the higher concentrations (Fig. 6B). The application of GA_3_ did not increase the fruit weight or change the fruit shape index of both genotypes (Fig. S8B-D).

**Figure 6:**
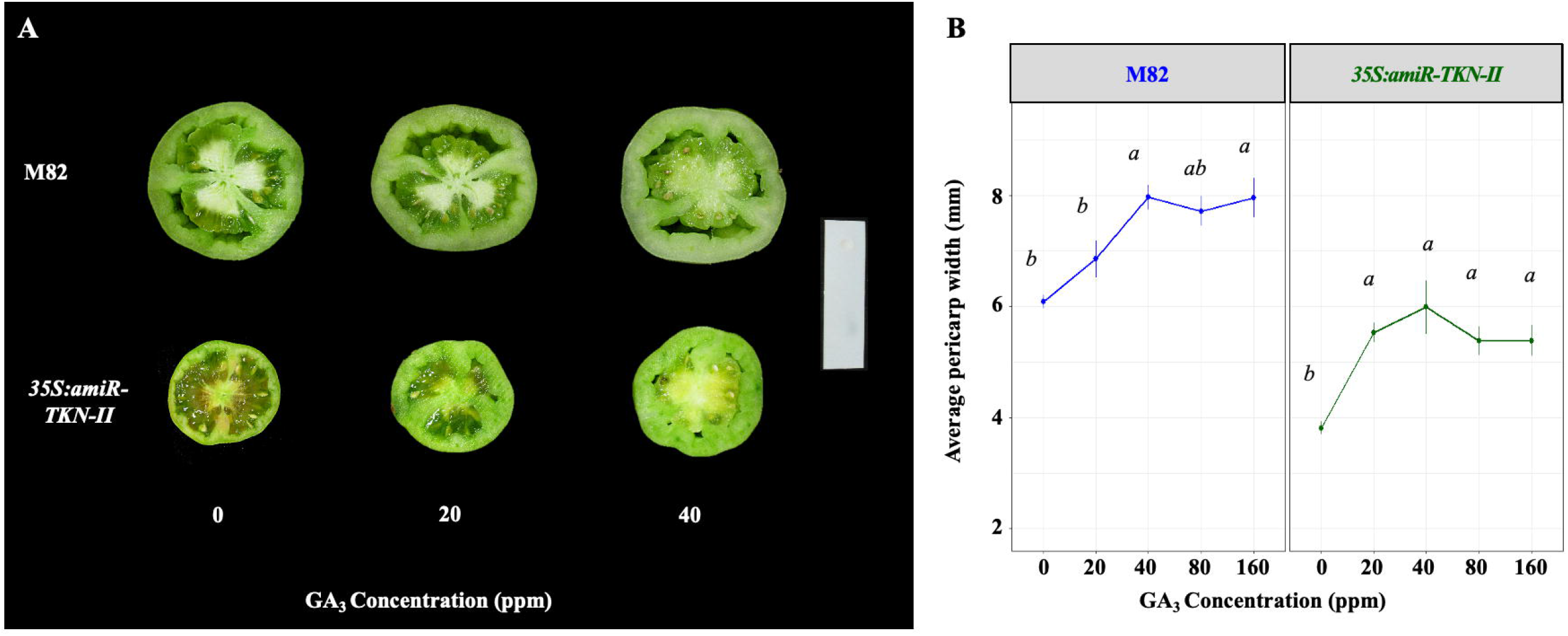
Ectopic GA_3_ application similarly affects M82 and *35S:amiR-TKN-II* pericarps expansion. **A:** Digitally extracted images of the 40 DAP fruit transverse sections from M82 line **top-panel** and *35S:amiR-TKN-II* line **bottom-panel**, treated with 0, 20, and 40 ppm of GA_3_ hormone during fruit development, show a correlation between the pericarp width and concentration of the GA_3_ treatment. The size bar = 6cm. **B:** Line plots displaying the average 40 DAP pericarp width (round dots) in mm (y-axis) of M82 (left panel) and *35S:amiR-TKN-II* (right panel) fruits treated with different concentrations of the GA_3_ (x-axis) show the pericarps expansion response to the GA_3_ application. The error bars represent standard errors. The number of sampled fruits for each genotype and treatment is ≥ 10. Different letters indicate Student’s-t-test H_0_ probability values ≤ 0.05, the similar letters indicate probability values > 0.05.

These results show that *35S:amiR-TKN-II* and M82 developing fruits respond similarly to external GA_3_ application, suggesting that elevated GA levels are insufficient to eliminate pericarp width and fruit size differences induced by the knockdown of the TKN-II genes.

## Discussion

### SlCLASS-II KNOX genes control fruit shape by regulating the orientation of cell divisions

Previous studies have established that ovary/pericarp abaxial-adaxial cell layers are mainly developed through the periclinal divisions of the pericarp layers. At the same time, cells along the ovary/fruit perimeter are a product of the anticlinal cell divisions (Renaudin et al., 2017). Examination of the pericarp anatomy phenotypes induced by the ubiquitous knockdown of the TKN-II genes detected fewer abaxial –adaxial cell layers and more E1 cells across the ovaries and pericarps of the knockdown fruits than in the M82 control. These observations suggest that TKN-II genes regulate the orientations and rates of cell divisions across the ovary/ fruit axis that determine the fruit shape (Wu et al., 2018). Current models suggest that tomato fruit shape is established during ovary development and controlled by the activity of the OVATE FAMILY PROTEINS (OFP) genes (Wu et al., 2018; Lazzaro et al., 2018), which were shown to interact with CLASS-II KNOX genes to control ovule development and secondary cell wall formation in *Arabidopsis thaliana* (Pagnussat et al., 2007; Li et al., 2011; Liu and Douglas 2015). Our results indicate that, similar to the *ofp* mutants (Wu et al., 2018), the shape index differences observed between M82 and *35S:amiR-TKN-II* mature fruits are established by the anthesis stage. These observations suggest that *TKN-II* / *SlOFP* interactions might play roles in tomato ovary development and the regulation of fruit shape.

### Reduced transcription levels of SlCLASS-II KNOX genes do not cause endoreduplication arrest

Examination of the *35S:amiR-TKN-II* fruit development has detected that the TKN-II genes promote pericarp cell expansion. Recent studies have shown that the expansion of pericarp cells is also promoted by the *GUANYLATE BINDING PROTEIN1* (*SlGBP1*) gene that controls the arrest of pericarp cells expansion and endoreduplication (Musseau et al., 2020). In the *TKN-II* knockdown fruits, the pericarp expansion was arrested at 30 DAP, while the expansion of the M82 pericarps continued until 40 DAP. However, unlike in *slgbp1* mutants, the early arrest of the *35S:amiR-TKN-II* pericarp expansion did not lead to an early arrest of the endoreduplication. These results suggest an uncoupling between endoreduplication and cell growth in *35S:amiR-TKN-II* pericarps, comparable to the previously reported observations in *ProPEPC2:SlKRP1* pericarps (Nafati et al., 2011).

### *SlCLASS-II KNOX* genes utilize GA-dependent and independent pathways to regulate fruit anatomy

Previous studies have established that the miss expression of CLASS-I KNOX genes in the leaves is reminiscent of the *class-II knox* mutant leaf phenotypes (Lincoln et al., 1994; Hareven et al., 1996; Janssen et al., 1998; Shani et al., 2009; Furumizu et al., 2015; Keren-Keiserman et al., 2022). That phenomenon is explained by the antagonistic functions of CLASS-I KNOX and CLASS-II KNOX genes in lateral organs (Furumizu et al., 2015).

The CLASS-I KNOX miss expression in tobacco, corn, and tomato leaves was shown to reduce GA levels by directly suppressing *GA20OX* (Sakamoto et al., 2001; Bolduc et al., 2012; Jasinski et al., 2008) or upregulation of the *GA2OX* (Bolduc and Hake 2009) gene expression. Moreover, the *SlTKN2* (CLASS-I KNOX) leaf miss-expression phenotypes were enhanced by the gibberellin deficient *gib2* mutation and suppressed by the *procera* mutant (Jasinski et al., 2008). These observations establish that CLASS-I KNOX genes modify lateral organ development via suppression of the GA activity (Jasinski et al., 2008) and suggest that CLASS-II KNOX genes regulate lateral organ development via the GA-mediated pathway.

Our examination of the *35S:amiR-TKN-II procera* fruit phenotypes suggests that *procera* leaf and ovary/fruit shape phenotypes are epistatic to the leaf and ovary/fruit shape phenotypes induced by the ubiquitous TKN-II knockdown. We also observed that *35S:PROΔ17* [constitutively suppressing GA activity (Nir et al., 2017)] and *35S:amiR-TKN-II* fruits had similar shape phenotypes. These results support the hypothesis that CLASS-II KNOX genes act via GA to regulate the tomato leaf and fruit shape (Fig. 7A).

**Figure 7:**
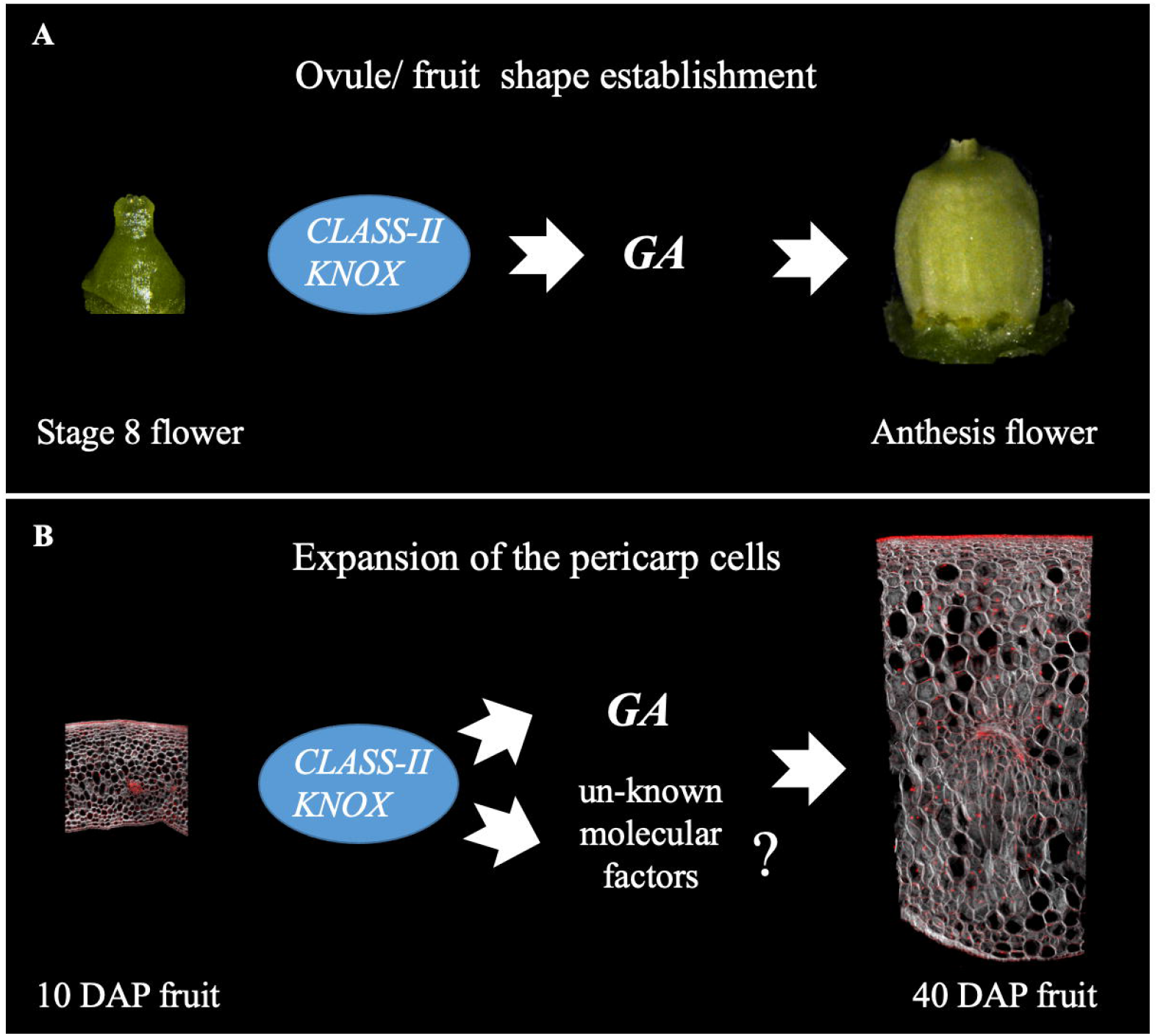
Tomato *CLASS-II KNOX* and Gibberellin mediated pathways interact in a developmental stage-dependent manner to control fruit morphogenesis. **A**-A model proposed for ovary/fruit shape establishment suggests that GA activity is epistatic to SlCLASS-II KNOX. **B-** A model proposed for pericarp cell expansion suggests the SlCLASS-II KNOX genes act via GA and additional unknown molecular factors to promote the expansion of the pericarp cells.

We also observed that the reduced pericarp expansion of the *35S:amiR-TKN-II* fruits is associated with elevated levels of the *SlGA2OX2* gene. However, the *procera* mutation failed to fully rescue the width, cell size, and ploidy levels of the *35S:amiR-TKN-II* pericarps. In addition, *35S:amiR-TKN-II* and M82 fruits responded similarly to ectopic GA application. These results suggest that TKN-II genes promote the expansion of the pericarp cells via both GA-dependent and independent pathways (Fig 7B).

### *CLASS-II KNOX* genes regulate fruit anatomy in multiple species

Previously reported silique phenotypes of the *Atknat3,4,5* triple mutants suggest that reduced AtCLASS-II KNOX activity results in shorter and broader pods with noticeably wider style (Furumizu et al., 2015 -Figure 3G; I; Furumizu et al., 2015 - Suppl. Figure 9). In *Arabidopsis*, the pod width is associated with mutations in genes controlling fruit shape (Roeder and Yanofsky, 2006; Wang et al., 2007; Wang et al., 2011), whereas the pod length is primarily controlled via the post-anthesis valve/ pericarp cell expansion (Vivian-Smith and Koltunow 1999; Ripoll et al., 2019). A recent analysis of the *Oryza sativa* kernels from the CRISPR-CAS9 generated *knat7* mutants, and *OsKNAT7* overexpression plants indicates that the gene regulates kernel shape and size (Wang et al., 2019). Combined with our observations, these results suggest that CLASS-II KNOX genes regulate fruit anatomy in multiple species and fruit types.

### Conclusions

The results presented here and in our recent report (Keren-Keiserman et al., 2022) advocate that during the development of tomato fruit, *SlCLASS-II KNOX* genes utilize diverse molecular and hormonal networks to regulate fruit anatomy and ripening.

## Accession Numbers

*SlKNATII3*: Solyc07g007120; *SlKNATII4:* Solyc12g010410; *SlKNATII5:* Solyc08g041820; *SlKNATII7:* Solyc08g080120; *SlGA20OX1*: Solyc03g006880; *SlGA20OX3*: Solyc11g072310; *PROCERA* Solyc11g011260; *SlGA2OX1*: Solyc02g070430; *SlGA2OX2*: Solyc07g056670; *SlGA2OX5*: Solyc10g007570

## Supplemental Data

**Table S1:** Gene abbreviations, genomic accession numbers, Forward and Reverse primers used for the RT-qPCR test. References to the previous studies using the same primers.

**Fig. S1:** Comparison of the equatorial perimeters and E1 cell length at the anthesis stage ovaries and 40 DAP M82 and *35S:amiR-TKN-II* fruits.

**Fig. S2:** Comparison of the abaxial –adaxial cell layer numbers in M82 and *35S:amiR-TKN-II* anthesis stage ovary wall, 10 and 40 DAP pericarps.

**Fig. S3:** Representative IMARIS reconstructed confocal images used for the histological analysis of the M82 and *35S:amiR-TKN-II* pericarps.

**Fig. S4:** Comparison of the GA metabolic, catabolic, and response regulators transcription levels in M82 and *35S:amiR-TKN-II* pericarps.

**Fig. S5:** Leaf and fruit phenotypes induced by the genetic manipulation of the GA response.

**Fig. S6:** Ovary wall and pericarp cytological phenotypes caused by the genetic interaction between the *pro*-loss-of-function mutant and TKN-II down-regulation.

**Fig. S7:** Box plots presenting ploidy distributions of the nuclei from the equatorial pericarps of 40 DAP -M82, *35S:amiR-TKN-II, procera (pro)*, and *35S:amiR-TKN-II procera* fruits.

**Fig. S8:** Phenotypes induced by the ectopic application of Gibberellin (GA_3_) to developing tomato fruits.

## Acknowledgments

We thank Yuval Eshed, Naomi Ori, Ilan Paran, and Idan Efroni for their valuable comments and manuscript discussion. We also thank Gil Yeshurun and Aharon Bellalou for their assistance at the greenhouse.

## Author contributions

A.G. supervised the research, A.G., N.G., E.F., and A.S. designed the experiments and analyzed the data. A.S., A.K-K., J.P-M., C.F, J.P.A., Z.A., N.G, E.M., and S.A-T., executed the experiments. A.G. wrote the manuscript.

## Data availability

All data is publicly available.

